# Development of methylation-based biomarkers for breast cancer detection by model training and validation in synthetic cell-free DNA

**DOI:** 10.1101/2022.02.11.480085

**Authors:** Sophie Marion de Procé, Martyna Adamowicz, Prasun Dutta, Sophie J. Warlow, Joshua Moss, Ruth Shemer, Yuval Dor, Christelle Robert, Timothy J. Aitman

**Author notes:** **Corresponding Authors** Prof. Timothy J. Aitman, Dr Christelle Robert. Usher Institute, University of Edinburgh Old Medical School, Teviot Place, Edinburgh, EH8 9AG (Sophie Marion de Procé).

## Abstract

Circulating tumour-derived DNA (ctDNA) carries the genetic and epigenetic characteristics of the tumour from which it is derived and can give information about the biology and tissue origins of the underlying tumour. DNA methylation is an epigenetic mark that is specific to individual tissues and, as methylation profiles are disrupted in tumours, they can indicate the tissue of origin and cancer type of ctDNA. We have developed a set of methylation biomarkers for detecting breast cancer in plasma cell-free DNA (cfDNA). First, we mined publicly available methylation datasets to create synthetic methylation profiles that were modelled to reflect cfDNA from healthy subjects and cancer patients. These profiles were restricted to the most differentially methylated CpGs between breast tumour samples and haematopoietic cells. Regularised logistic regression models were trained using 10-fold cross-validation on synthetic cfDNA datasets with distinct fractions of breast tumour DNA spiked *in silico* into healthy cfDNA with the addition of 10% of a mix of different tissues. Initial validation with synthetic cfDNA permitted detection of breast cancer-derived DNA with as little as 0.25% tumour DNA spiked *in silico* into healthy subject cfDNA with an area under ROC curve (AUC) of 0.63. Performances of classifiers increased with increased fractions of spike-in tumour DNA (AUCs 0.77 and 0.93 at tumour DNA fractions 0.5% and 1% respectively). We then combined the most discriminative CpG markers from our models with methylation markers of breast cancer that had already been published to obtain a single marker set. *In vitro* testing of MCF-7 breast cancer cell line DNA spiked into leukocyte DNA showed highly significant correlation for individual markers between laboratory-measured and published methylation data for MCF-7 and leukocytes (R > 0.89, *P* < 2.2 × 10^−16^). These preliminary data indicate promising results for detection of breast cancer cell line DNA using this methylation marker set, which now require testing in cfDNA from breast cancer patients and healthy controls.

## Introduction

Cancer is a leading cause of death, accounting for nearly 10 million deaths worldwide in 2020 (Sung et al., 2021). Female breast cancer is the most prevalent cancer, with 2.3 million women diagnosed with breast cancer and 685,000 deaths globally in 2020, with breast cancer surpassing lung cancer as the most commonly diagnosed cancer (Sung et al., 2021). A key challenge in reducing mortality is to detect the disease and any potential residual disease or recurrence as early as possible. Early detection and treatment is widely viewed as being central to improving patient survival (Hawkes, 2019; CRUK, 2020).

The last decade has seen steadily increasing interest in the potential of liquid biopsy, through the analysis of plasma cell-free DNA (cfDNA), to detect diseases such as cancer (Corcoran and Chabner, 2018; Keller et al., 2021). Indeed, a non-invasive, sensitive, specific, and readily repeatable test would be of significant clinical value. Much work has been undertaken aiming to detect tumour-derived mutations in DNA that has been released from tumour cells into the circulation, the fraction of cfDNA that is known as circulating tumour DNA (ctDNA). cfDNA is found at low concentrations in plasma with a short half-life of less than an hour and, in patients with small or early-stage cancer, ctDNA comprises a small, often tiny fraction of total circulating cfDNA. Of importance in cancer management, ctDNA retains the genetic and epigenetic characteristics of the tissue from which it was derived, including the carriage of somatic mutations and tissue-specific methylation profile.

Different tissues and cell types have consistent genome-wide methylation profiles although such profiles show some inter-individual variation. In addition, since tumours have disrupted methylation patterns, the analysis of methylation patterns of plasma cfDNA can permit identification of tumour-derived DNA in patients with cancer, as well as the tissue(s) of origin of an individual’s circulating cfDNA. Several groups have already investigated methylation in cfDNA to detect its tissue-of-origin, with generally encouraging results (Moss et al. 2020; Liu et al. 2020).

cfDNA in healthy people originates mostly from the blood cell lineage, comprising around 55% leukocyte-derived DNA, and 30% from erythroblast DNA with additional smaller contributions from other tissues (Lam et al. 2017; Moss et al. 2018). In this report, we aimed to design a set of CpG biomarkers that could reliably detect breast cancer DNA in liquid biopsy samples by mining of public DNA methylation databases and by searching for markers of breast cancer that are already published.

## Materials and Methods

### Public data download and processing

All analyses were performed using the R statistical package (Core Development Team, 2020). Global methylation profiles from biological samples hybridised onto the Illumina Infinium HumanMethylation450K BeadChip were downloaded from The Cancer Genome Atlas (TCGA) using the TCGAbiolinks R package (Colaprico et al., 2016; Silva et al., 2016; Mounir et al., 2019) and from Gene Expression Omnibus (GEO) (Edgar et al., 2002; Barrett et al., 2013) (Table 1). The idat binary files were pre-processed together using the subset-quantile within array normalization (SWAN) method, mapped to the hg19 genome, and ratio-converted into β-values (ratio of methylated to unmethylated cytosines in the target CpG motif) and M-values (the logarithmic transformation of β-values) using the Bioconductor minfi package (version 1.22.1) (Maksimovic et al., 2012; Aryee et al., 2014). Probes containing a SNP at the CpG interrogation and/or at the single nucleotide extension, for any minor allele frequency were removed from the dataset.

**Table 1.** DNA methylation samples (450K methylation array) from published TCGA and GEO (Demetriou et al., 2013; de Goede et al., 2015) datasets used in this study.

| **Tissue** | **Neoplastic Status** | **Source** | **Number of samples** |
| --- | --- | --- | --- |
| breast | cancer | TCGA-BRCA | 793 |
| breast | normal | TCGA-BRCA | 97 |
| lung | normal | TCGA-LUAD | 32 |
| liver | normal | TCGA-LIHC | 50 |
| kidney | normal | TCGA-KIRC | 160 |
| colon | normal | TCGA-COAD | 38 |
| pancreas | normal | TCGA-PAAD | 10 |
| leukocytes | normal | GSE51057 | 329 |
| erythroblasts | normal | GSE68456 | 12 |
| erythroblasts | normal | GSE82084 | 9 |

### Differential methylation analysis

We chose to restrict synthetic cfDNA assembly to the most differentially methylated features in the 450K methylation array. We performed differential methylation analysis by comparing methylation profiles from haematopoietic cells (leukocytes and erythroblasts, n=350) (Demetriou et al., 2013; de Goede et al., 2015) with breast tumour samples (n=793) using the dmpFinder function of the minfi package. The 450K array features were ranked by decreasing q-value and then by the difference of the medians of the β-values and the top 3000 features were retained. This follows the widely used method of comparing tumour samples to leukocytes to detect cancer using liquid biopsy (Lehmann-Werman et al., 2016).

### *In silico* generation of synthetic cfDNA

We assembled synthetic methylation profiles simulating cfDNA assuming that the origin of healthy cfDNA is mostly haematopoietic DNA, with minor contributions from other tissue types (Figure 1). By further compounding healthy synthetic cfDNA samples with a range of fractions of tumour methylation profiles, we aimed to set up an *in silico* detection test for cancer DNA in liquid biopsies, by training a model on methylation features to discriminate between healthy and cancer synthetic cfDNA profiles. Synthetic cfDNA samples were derived as the result of mixing healthy and cancer β-value profiles according to pre-defined proportions, or mixture coefficients. Samples were mixed by multiplying each sample vector of β-values by its mixture coefficient, and then adding up the resulting CpG methylation values.

**Figure 1:**
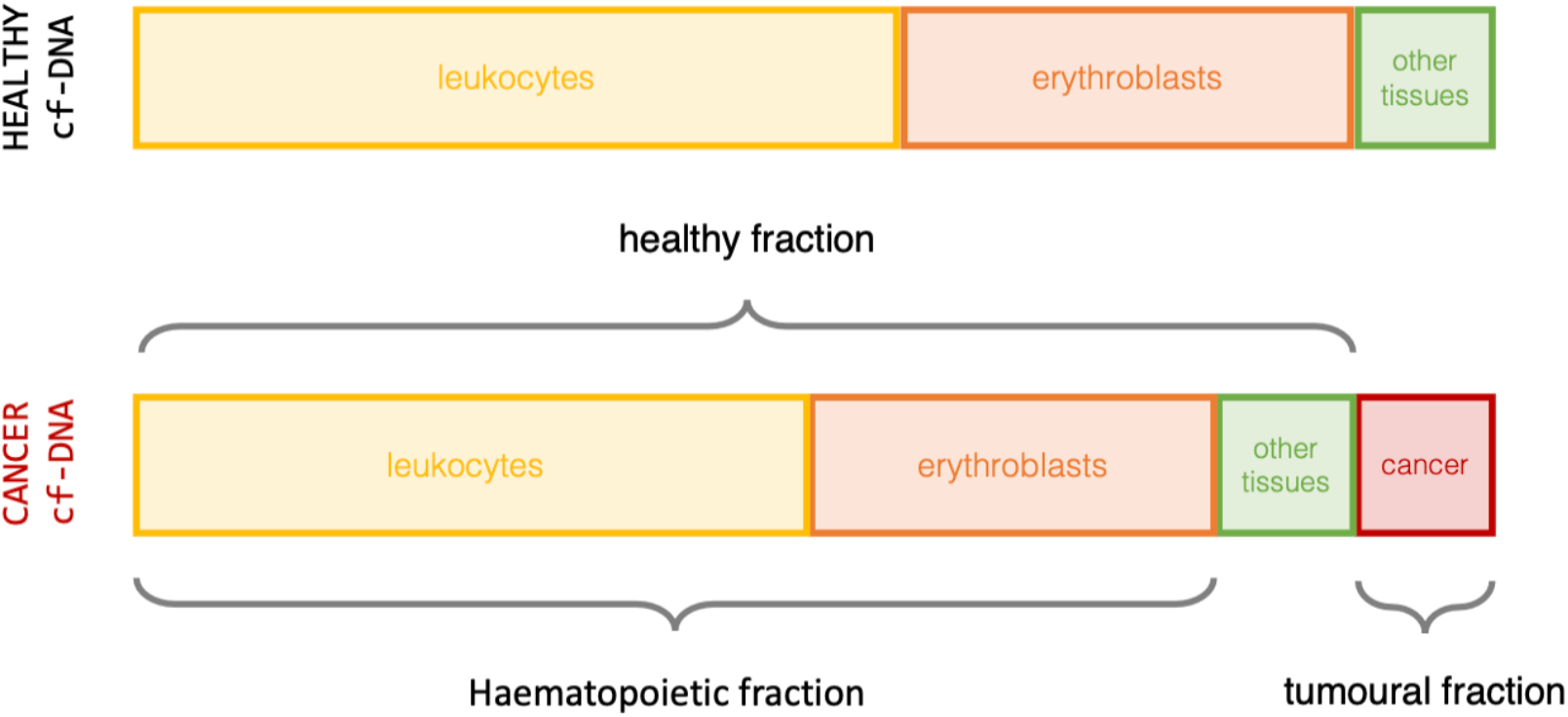
Synthetic cfDNA construction. Synthetic cfDNA samples were assembled by mixing global methylation profiles of different tissue types in controlled proportions. First, haematopoietic synthetic profiles were created by mixing a single leukocyte and a single erythroblast profile in a 70:30 ratio. Then, a non-haematopoietic “other tissues” profile (a single random healthy profile of each of breast, lung, liver, kidney, colon or pancreas) was added to make up 10% of the resulting healthy cfDNA synthetic profile. Finally, cancer synthetic cfDNA profiles were the result of mixing a single cancer methylation profile, in a pre-defined fraction, to a healthy profile.

Haematopoietic profiles were first assembled by mixing each of the 329 available leukocyte profiles with a random erythroblast profile, in a 70:30 ratio. Then, a single random healthy profile from breast, colon, liver, kidney, lung or pancreas (with equal probability of selection across the six tissue types), was mixed with the haematopoietic profile, in 90:10 ratio, yielding a set of 329 healthy synthetic cfDNA profiles. Another set of 329 healthy cfDNA samples was constructed in the same way and then mixed with a single randomly sampled breast tumour methylation profile, in a set of tumour DNA proportions of 0.01%, 0.1%, 0.25%, 0.5%, 0.75%, 1% and 10%. This resulted in 7 different hypotheses tested on healthy and cancer-derived cfDNA. For each scenario, there were thus 329 simulated healthy cfDNA profiles, and 329 simulated cancer patient cfDNA profiles (Figure 1). The construction of the 658 cfDNA samples was repeated to obtain 10 replicates for each percentage of tumour DNA, using the same random profiles for each replicate across the 7 percentages of tumour DNA tested.

### Model training and testing on synthetic cfDNA

Seventy models (10 for each tumour DNA proportion) were trained on methylation profiles of synthetic cfDNA samples to classify cfDNA as coming from either breast cancer patients or healthy people. Each set of 329 cfDNA samples was divided into a training set with approximately 80% of the data and a testing set with the remaining 20%. There are therefore 263 healthy cfDNA and 263 cancer cfDNA profiles in the training set, and 66 healthy cfDNA and 66 cancer cfDNA profiles in the testing set.

We trained generalised linear models on the training set via penalised maximum likelihood with 10-fold cross-validation provided in the glmnet R package (Friedman et al., 2010). L1 penalised logistic regression provides an efficient lasso regularization path for logistic regression, which results in models with less features and smaller coefficients. The regularization path was computed for the lasso penalty for a range of values for the regularization parameter lambda. The models corresponding to the largest value of lambda such that error is within 1 standard deviation of the minimum were then retained and used to classify synthetic cfDNA samples.

Each of the 70 models was used to classify cancer cfDNA from the test data set. We computed the area under the receiver operating characteristic (ROC) curve (AUC) as a metric for the classification performance of the models. For each given tumour fraction, ten AUCs were computed based on ten sets of 329 synthetic cfDNA samples assembled independently (Table 2, Supplementary Figure 1).

**Table 2.** Performances of linear models based on synthetic cell-free DNA methylation profiles for distinct percentages of breast tumour DNA. A total of 70 linear models were built using our *in silico* classification analysis accounting for 7 different percentages of breast tumour DNA and 10 replicates. Mean area under the ROC curve (AUC) and standard deviation were calculated across 10 replicates per percentage of tumour DNA. Performances of each of the 70 classifiers along with the number of features (CpGs) selected are available in Supplementary Table 2.

| **Percentage of tumour DNA** | **Mean AUC** | **Standard Deviation** |
| --- | --- | --- |
| 0.01 | 0.508 | 0.007 |
| 0.1 | 0.526 | 0.034 |
| 0.25 | 0.628 | 0.001 |
| 0.5 | 0.771 | 0.013 |
| 0.75 | 0.873 | 0.004 |
| 1 | 0.931 | 0.01 |
| 10 | 1 | 0 |

To obtain a set of CpGs to test on physiological cfDNA, we counted for each CpG the number of models in which that CpG had a non-zero coefficient, and then ranked the CpGs to pick the CpGs appearing in the largest number of models. Since each model has 10 replicates, the maximum number of models a CpG could be found in was 70. This method assumes that CpGs that are selected by glmnet in many different models are likely to be good discriminants of cancer in cfDNA. We selected the 100 CpGs that were found in the most models for experimental analysis.

### Methylation biomarkers from the literature

We selected papers describing methylation biomarkers to detect breast cancer. We searched recent scientific articles mentioning breast cancer methylation biomarkers and identified 11 articles with methylation biomarkers of potential interest. Out of these, 3 articles had listed CpGs from the Illumina Infinium Methylation 450 arrays (Uehiro et al., 2016; de Almeida et al., 2019; Fackler et al., 2020). From de Almeida et al.’s study (de Almeida et al., 2019), we selected the top 55 CpGs as ranked by differential methylation p-value. From Fackler et al.’s study (Fackler et al., 2020), we selected their panel of 30 CpGs with β < 0.01 in all normal breast samples, as well as β > 0.015 in at least 4 of 110 tumours. From Uehiro et al.’s study (Uehiro et al., 2016), we used all 12 loci. In addition, 13 CpGs were provided by the Moss group on a collaborative basis (Moss et al. 2020). We used the markers derived from these sources to compile a list of 110 CpGs in our experimental assay that were added to the markers derived from data mining and modelling to create a panel of 210 marker CpGs (Supplementary Table 1). There were no overlapping CpGs between the literature-derived markers and those derived from our data mining and model training.

### Input DNA for laboratory in-house developed targeted methylation assay

Commercial gDNA was used for the assay validation: MCF-7 cell line (Merck, #86012803), leukocyte (AMS Bio, #D1234148) and CpG Methylated HeLa (NEB, #N4007). The RepliG mini kit (Qiagen, #150023) was used to generate fully non-methylated HeLa DNA. 1 µg of DNA per sample was sheared to the average fragment size of 160 bp using a Covaris S220 sonicator (Covaris, #500217). DNA quality was validated using the TapeStation 4200 system (Agilent, #G2991BA) and Genomic DNA ScreenTape (Agilent, #5067-5365). DNA quantity was estimated using Qubit fluorometer (Invitrogen, #Q33226) and Qubit dsDNA BR Assay (ThermoFisher, #Q32850). 20 ng of sheared DNA was used for library preparation. MCF-7 spike-in samples were prepared by adding MCF-7 DNA to leukocyte DNA at 1%, 5%, 12.5%, and 25%.

### Library preparation and sequencing

20 ng of sheared DNA was used to prepare target enrichment libraries incorporating error suppression including unique molecular indexes (UMIs) and unique dual indexes (UDIs) which remove PCR and sequencing errors as well as index hopping events, providing uniformity of coverage. DNA fragments were end-repaired and A-tailed using Cell3™ Target: DNA Target Enrichment kit (Nonacus Ltd, #C3212RK), dual methylated adapters including UMIs were ligated to repaired fragments. The constructs were subjected to treatment with NEBNext® Enzymatic Methyl-seq (EM-seq™) (NEB, #E7125) according to the manufacturer’s protocol. The method provides an enzyme-based alternative to bisulfite conversion. Following amplification using Cell3™ Target: DNA Target Enrichment kit and NEBNext® Q5U® Master Mix (NEB, # M0597), DNA quality and quantity were measured using TapeStation 4200 system, D1000 ScreenTape (Agilent, # 5067-5582), Qubit fluorometer and Qubit dsDNA BR Assay Kit (ThermoFisher, #Q32850), individual libraries were pooled to a total amount of 1 μg and enriched using Cell3 ™ Target Hybridization and Capture Kit (Nonacus Ltd, #C3CUST) and double-stranded DNA baits, in-house developed targeted methylation assay, to target 210 CpG biomarkers. The quality of the final libraries was validated using the TapeStation 4200 system, High Sensitivity D1000 ScreenTape (Agilent, # 5067-5584), Qubit fluorometer and Qubit dsDNA HS Assay (ThermoFisher, # Q32851). Finally, libraries were sequenced using MiSeq (Illumina), generating 150 bp paired-end reads.

### Sequence analysis

Raw BCL (base call files) generated using Illumina MiSeq instrument were downloaded from BaseSpace which were then processed using bcl2fastq v2.17.10 to generate paired-end fastq files for each sample. During BCL to fastq conversion procedure, the RunInfo.xml file was edited to obtain an additional fastq file for sequenced UMI codes associated with each read, resulting to a total of three fastq files per sample (Read 1, Read 2 and UMI sequence).

Each read name of the paired-end fastq file was associated with its corresponding UMI sequence to create new set of fastq files using a custom script. The UMI was attached to be the last element of the read name. Quality control visualisation was performed using FASTQC v0.11.9. TrimGalore v0.4.1 was used with parameters --paired -q 30 --clip_R1 10 --clip_R2 20 for removing adapters, low quality bases from 5’ end of the reads. Trimmed paired-end fastq files were aligned to the converted versions (C->T and G->A converted) of the human genome (GRCh38) using bismark v0.22.3 with -- pbat (align reads only to complementary to (converted) top strand and bottom strands) and --local options. UMI based deduplication was performed on the resulting BAM file using deduplicate_bismark v0.22.3 script with –paired and –barcode options. Finally, the bismark_methylation_extractor v0.22.3 script was used to extract methylation values for every single C analysed. The output files were converted to bedGraph form using bismark2bedGraph v0.22.3. The bedGraph files were further processed using custom script to obtain methylation values for both forward and reverse strands for all sequenced CpGs.

To generate methylation percentage values per target region of interest, a custom Perl script was written which utilised a bed file containing coordinate information of targeted CpGs in the marker set +/-100 bp along with the bedGraph file produced above. This script in turn utilised a custom R script that performed the actual methylation % value calculation. Methylation percentage was calculated for each CpG site present in the target region which had >= 5 reads and an average was calculated of the resulting values to calculate methylation percentage per target region. Further downstream analysis and visualisation was performed using custom R scripts.

### Comparison of *in vitro* data with 450k array

The methylation percentage values of CpG sites from *in silico* modelling were compared to their expected pattern seen in a 450K methylation array. This was done by calculating the correlation between *in vitro* methylation percentage of our CpG sites of interest and methylation percentage (calculated from beta values) of the same sites in the 450K array. Pure MCF-7 control cell line idat files were downloaded from GEO with accession number: GSM2492223. Leukocyte idat files, containing methylation profile of peripheral leukocytes from 117 healthy controls, were downloaded from GEO with accession number: GSE67393. Both sets of data were processed using the minfi R package (Aryee et al., 2014).

Briefly, for MCF-7, the idat files were first pre-processed and normalised using the ssNoob (single sample normal-exponential using out-of-band probes) method (Fortin et al., 2017) from the minfi package. Before normalisation, a check was done to ensure that the mean detection p-value for the sample was > 0.01. Probe level filtering was performed whereby failed probes with detection p-value < 0.01 were filtered out and probes present on sex chromosomes were also filtered out to remove sex-based effects on methylation profile. Further probe filtering was performed as recommended by Zhou et al. (Zhou et al., 2017). Specifically, those probes were removed which were marked TRUE for the MASK_GENERAL (recommended general purpose masking) column in the probe annotation file provided by Zhou et al. (https://zhouserver.research.chop.edu/InfiniumAnnotation/20180909/HM450/HM450.hg38.manifest.tsv.gz). Beta values were then obtained for those probes whose target CpG sites were common to our set under study.

The exact same procedure was followed to pre-process leukocyte samples. In addition to probe filtering, sample filtering was performed where samples were removed which had median intensity < 10.5. Since there were multiple samples in the dataset, the mean was calculated for all the beta values across the samples for each CpG locus. Using the beta values from MCF-7 and leukocyte dataset, *in silico* MCF-7 and leukocyte spike-in was created which was then compared with the *in vitro* data by calculating the correlation coefficient. Beta values were multiplied by 100 to get the actual methylation percentage value before correlation calculation. All analysis were done using custom scripts in R.

## Results

We used an *in silico* modelling approach to build 70 linear models to classify breast cancer patient cfDNA from healthy people cfDNA for a range of 7 breast tumour percentages using synthetic methylation profiles (Table 2, Supplementary Table 2). This approach shows that, as expected, the areas under the ROC curve (AUC) increase as the percentage of tumour DNA in the cfDNA increases. We can classify between synthetic cfDNA from healthy people and synthetic cfDNA from cancer patients when there is as little as 0.25% of tumour DNA in the cfDNA with mean AUC of 0.63 (Table 2). For increased fractions of breast tumour DNA in cfDNA from 0.5% to 1%, classification performance increases with mean AUC of 0.77 to 0.93 respectively. At 10% of spike-in breast tumour DNA, the models show the highest classification performances (AUC of 1) with perfect discrimination between tumour and healthy cfDNA samples. Examples of ROC curves for the models trained to classify synthetic cfDNA into healthy or breast cancer at seven different percentages of breast tumour DNA are shown in Supplementary Figure 1.

We selected the 100 CpGs that were a feature in the highest numbers of models as a proxy for good classification power. A set of these 100 CpGs from our *in silico* modelling approach and 110 CpGs from our literature search was tested on our experimental assay on the breast cancer cell line MCF-7 DNA, normal breast DNA and serial dilutions of MCF-7 into leukocyte DNA.

We generated sequencing libraries for methylated and non-methylated control DNAs, and for, MCF-7 cell line DNA with 1%, 5%, 12.5%, and 25% MCF-7 spike-in on a leukocyte background. We obtained 120-450 ng per pre-capture library yielding enough to pool 1 μg of combined DNA for sequencing. Following enrichment and amplification a final library was generated with average fragment size 330 bp. Library sequencing and demultiplexing yielded a median of 798,700 paired-end reads (range 601,685 - 1,300,303) per sample; a median coverage of 504 x per sample (range 325 - 704 x) was obtained, with 0.9 – 2.2 million total mapped reads per sample (Supplementary Tables 3 and 4).

Looking for evidence of our assay’s potential to discriminate between samples from breast cancer patients and samples from healthy people, we plotted heatmaps of the methylation percentage values for all CpGs together (Figure 2) and for CpGs from *in silico* modelling and literature separately (Figure 3). The control DNA samples indicated a high conversion rate for the enzymatic conversion protocol, with 0.2% methylation called on RepliG-amplified unmethylated control DNA and 96% methylation percentage on HeLa DNA. In addition, a large proportion of markers from both sets of CpGs are indicative of the percentage of MCF-7 and leukocyte DNA (Figure 3). Similarly, when markers derived by data mining and modelling are separated into those expected to show hypermethylation (hyperM) or hypomethylation (hypoM) in breast cancer compared to leukocytes, marked differences were seen between samples according to percentage of MCF-7 and leukocyte DNA (Figure 4), with differences visible from the smallest percentages of MCF-7 between 1% and 5%. The differences between samples can be clearly observed in correlation score heatmaps (Supplementary Figure 2). When looking only at the CpGs from the *in silico* modelling, we can compare the methylation data generated *in vitro* with the expected pattern of these CpGs from 450K data. HyperM CpGs should have high methylation in breast tumour and low methylation in leukocytes, with the converse expected for HypoM CpGs. This is well reflected in our experimental data, with highly significant correlation for individual CpGs between the *in vitro* data at 100% MCF-7 DNA and 1% MCF-7 DNA and 450K data for these markers in published MCF-7 and leukocyte 450K array data (Figure 5).

**Figure 2:**
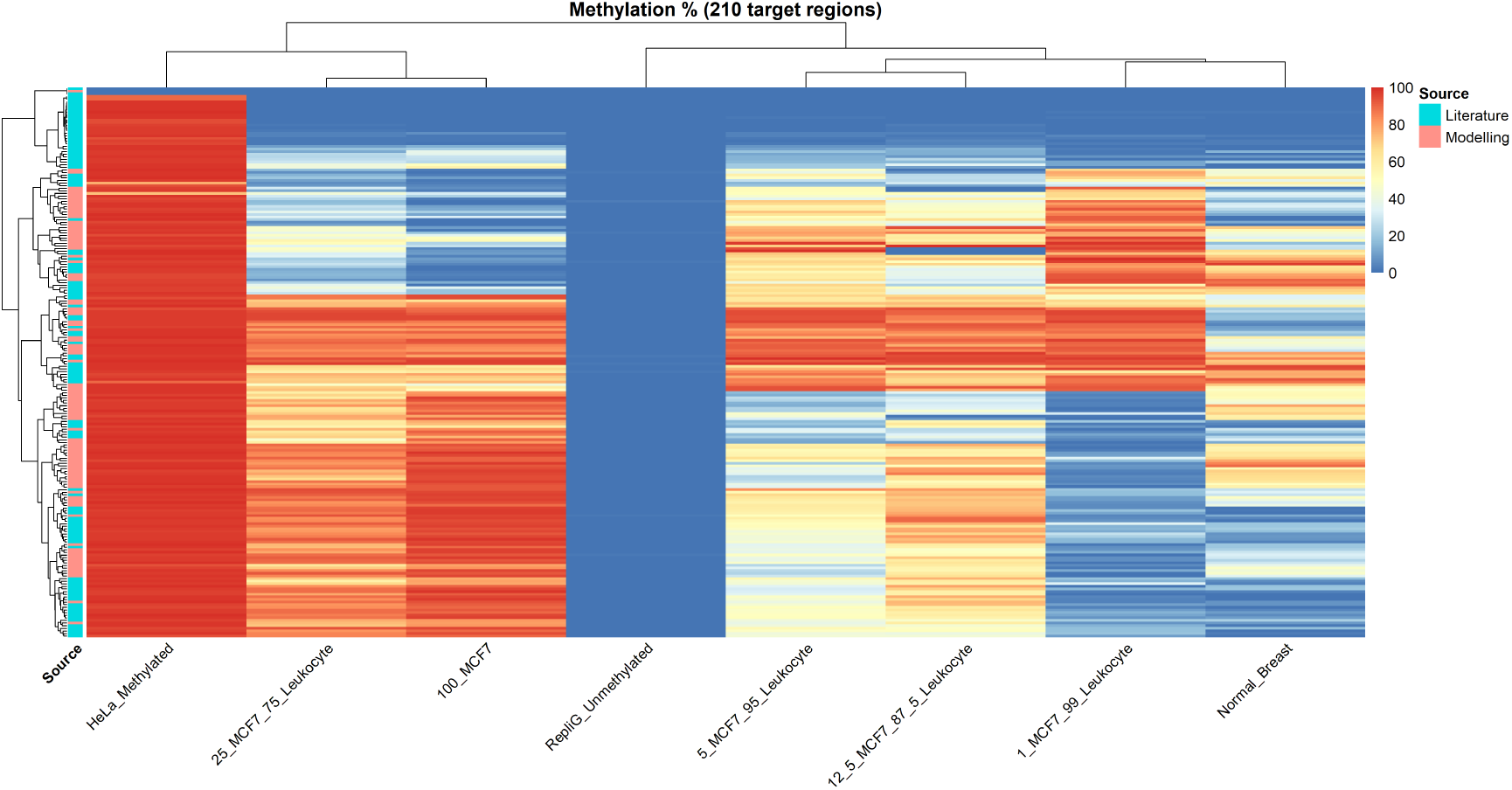
Heatmap showing methylation percentage values, determined by our in vitro methylation assay, of 210 selected target regions containing our CpGs of interest. The experiment used MCF-7 DNA spiked with different percentages of leukocyte DNA (99%, 95%, 87.5%, and 75%) along with methylated (HeLa_Methylated) and unmethylated (RepliG_Unmethylated) control samples and a pure MCF-7 sample (100_MCF7). CpG markers were derived from both the data mining/modelling methods and the published literature. Hierarchical clustering of methylation percentage values of the target region containing CpGs of interest revealed clear distinction between different samples. Control samples (fully methylated and unmethylated) showed high and low methylation percentage values, respectively.

**Figure 3:**
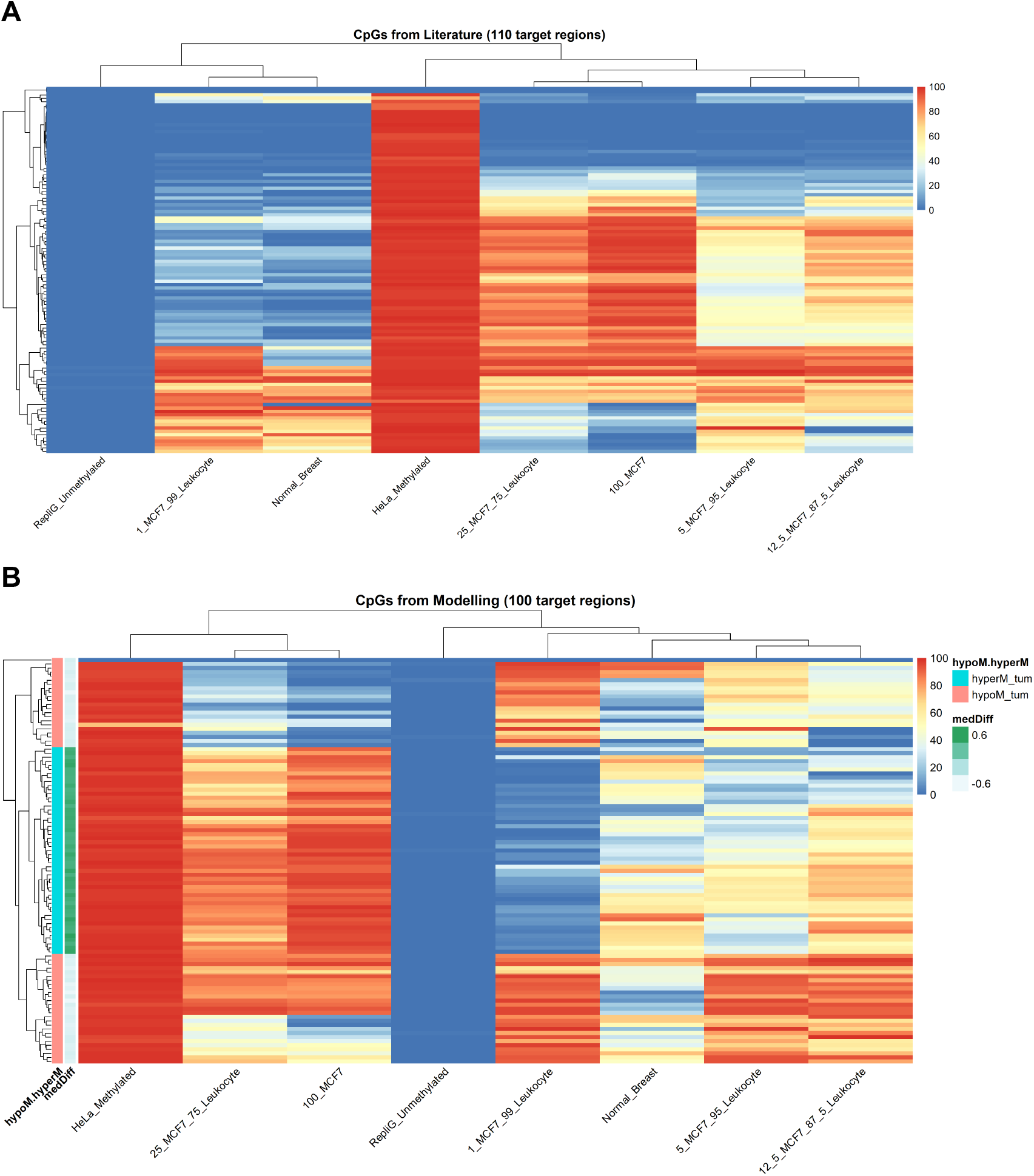
(A) Heatmap showing methylation percentage values, determined by our in vitro methylation assay, of 110 selected target regions containing our CpG of interest sourced from literature. (B) Heatmap showing methylation percentage values, determined by our in vitro methylation assay, of 100 selected target regions containing our CpG of interest sourced from data mining and modelling. Additional annotation provides information about if the target region is hypo or hyper methylated (HypoM_tum and HyperM_tum respectively) in breast cancer in the sourced data compared to haematopoietic cell DNA. HyperM_tum indicates markers with breast tumour methylation higher than leukocytes and converse in HypoM_tum. medDiff is the median difference in methylation percentage: positive numbers indicate breast tumour markers with higher methylation compared with Buffy coat. HyperM_tum and HypoM_tum CpGs form two main clusters which clearly correlate with medDiff values.

**Figure 4:**
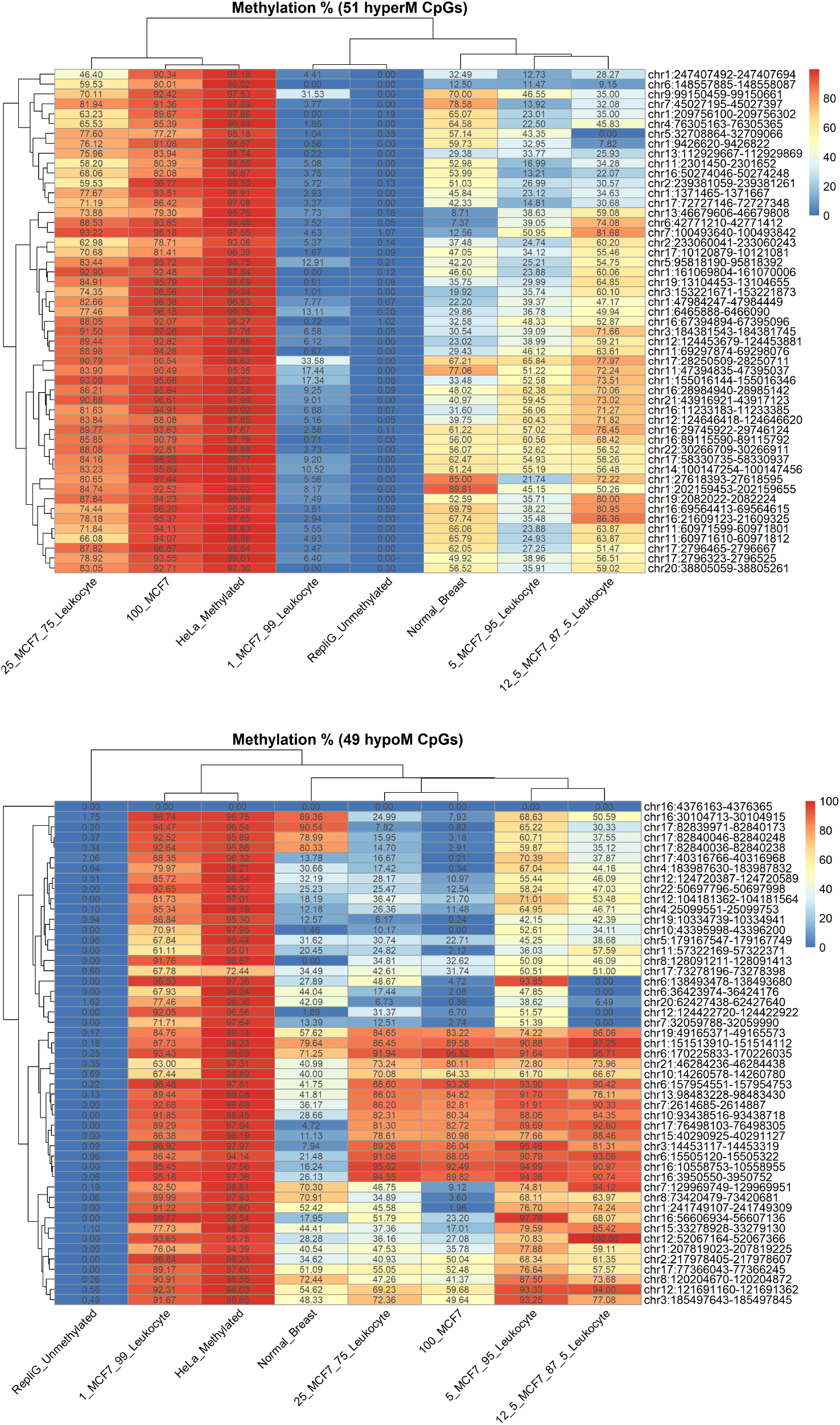
Heatmap of methylation percentage values of 100 CpG target regions, determined by our in vitro methylation assay, that were expected to be hypermethylated (hyperM; Upper Panel) or hypomethylated (hypoM; Lower Panel) in breast tumour compared to haematopoietic cells. Hela and RepliG control samples show respectively very high and low methylation percentage values. 100_MCF7 sample are distinct from Normal_Breast sample. In hyperM CpGs, methylation percentage decreases as MCF-7 percentage decreases and leukocyte percentage increases in sample mixtures. In hypoM CpGs, methylation percentage for most markers decreases with increasing percentage of MCF-7. The genomic coordinates of each target region are shown to the right of the heatmap.

**Figure 5:**
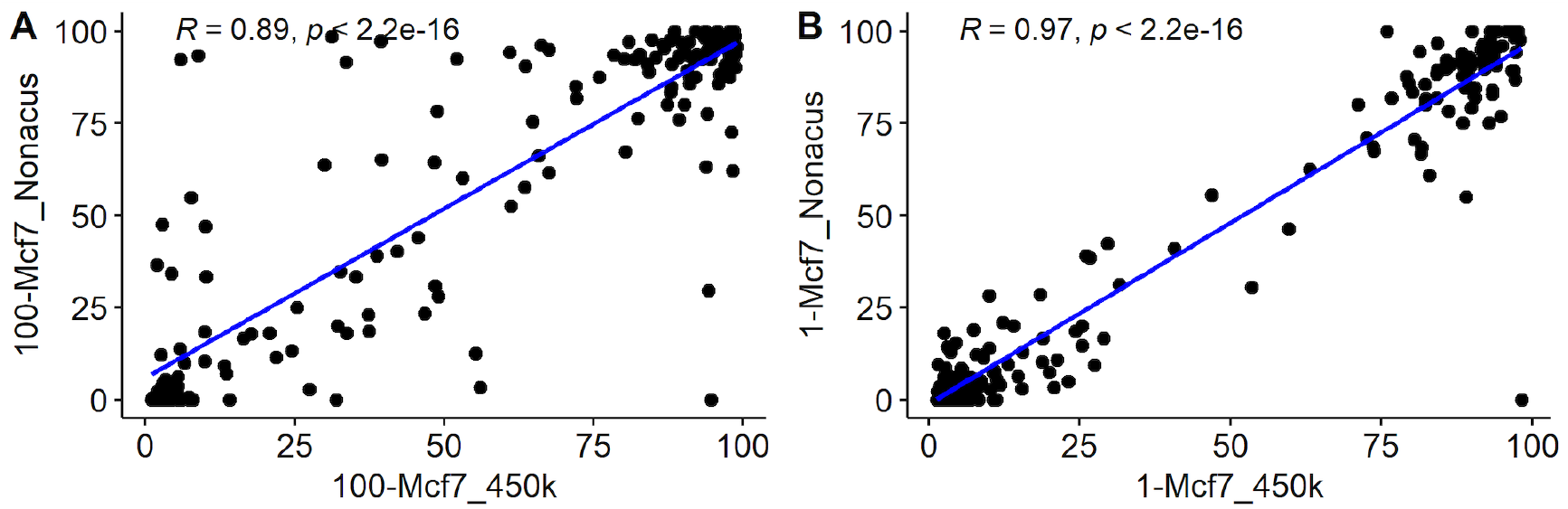
Scatter plots showing the correlation of individual marker CpG methylation percentage for pure MCF-7 DNA between *in vitro* generated data in this study and published 450K data for MCF-7 (A) and for the *in vitro* mixture of 1% MCF-7 with 99% leukocyte DNA and *in silico* modelled 450K MCF-7 and leukocyte methylation profile with the same tissue proportions (B). The blue line represents the line of best linear fit, with Pearson correlation coefficient and p-value shown.

We anticipated that, in the data we generated *in vitro*, the CpGs with the highest and most significant correlations between measured methylation percentage and the percentage of MCF-7 DNA in each sample will have the best power to distinguish between cfDNA from healthy people and cfDNA from breast cancer patients. A correlation analysis of the measured methylation percentage against percentage of MCF-7 DNA spiked into leukocyte DNA suggests that most CpGs have a high absolute value of correlation coefficient, for which the six most significant correlations are shown in Figure 6. In addition, when comparing the correlation coefficients of markers derived from modelling and those derived from the literature, there was no significant difference in the absolute value of correlation coefficients or p-values between the CpGs from either source (Figure 7, p>0.6). The subset of CpGs associated with numerically low correlation coefficients in both modelling and literature-derived marker sets might indicate MCF-7 specific CpGs with MCF-7 cell lines representing only a subset of the heterogeneity of breast cancer methylation profiles.

**Figure 6:**
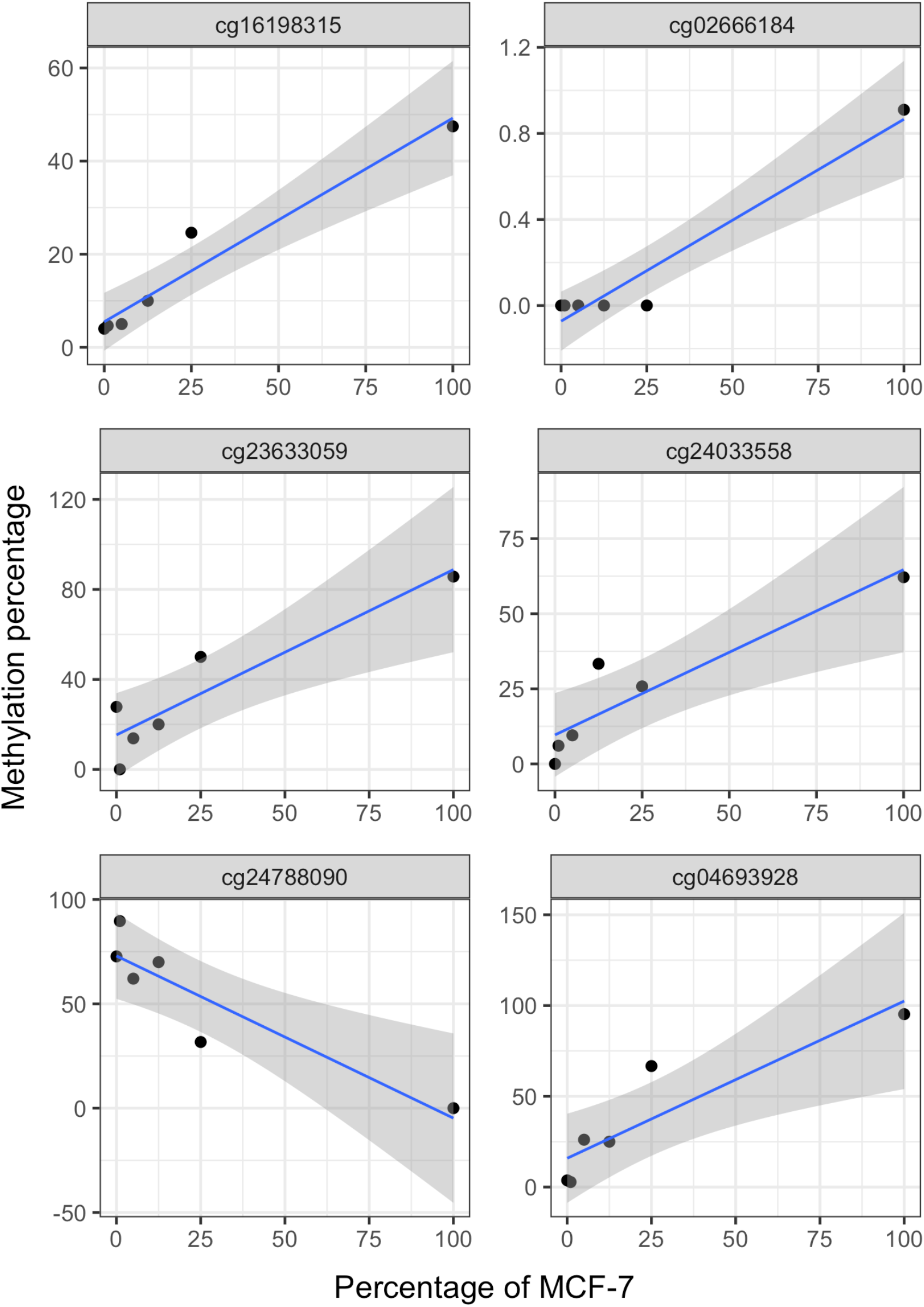
Methylation percentage from our in-house developed targeted methylation assay plotted against percentage of MCF-7 DNA spiked into leukocyte DNA for each of the 6 CpGs with the lowest p-value for Pearson’s product-moment correlation test. The blue line represents the regression line fitted to the data using a linear model. The grey area represents the 95% confidence level interval for predictions from a linear model.

**Figure 7:**
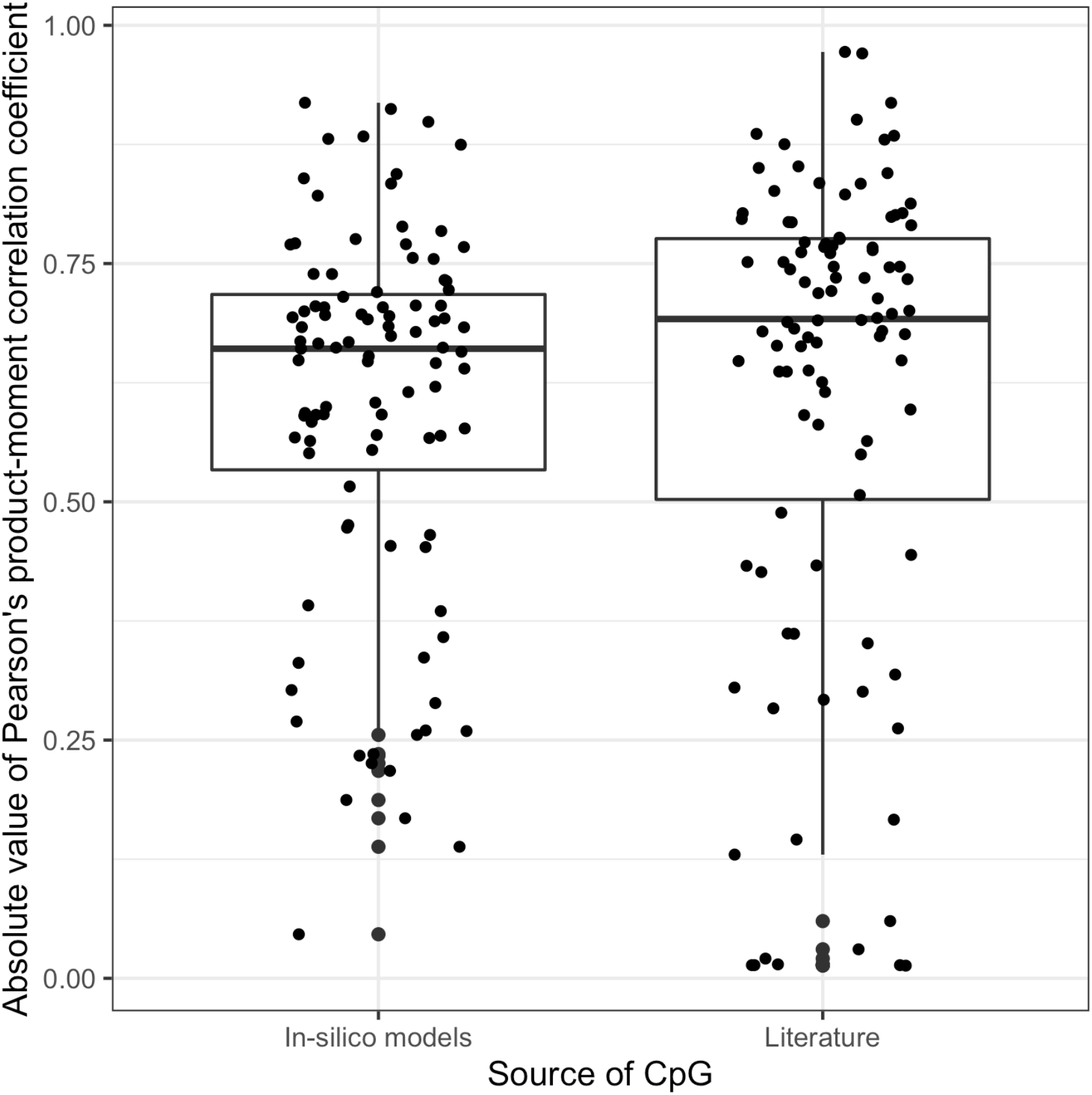
Boxplot of the absolute value of the Pearson’s product moment correlation coefficients between methylation percentage measured *in vitro* and percentage of MCF-7 in each sample, for each source of CpGs in the methylation assay.

## Discussion

Our *in silico* modelling showed that the markers we derived by data mining of publicly available data were able to detect the presence of breast cancer DNA when this DNA contributes to as little as 0.25% of synthetically modelled cfDNA. This is encouraging but remains to be tested on healthy people and breast cancer patient cell-free DNA samples, as the 450K array data will have different properties compared with our assay’s sequencing data, and the modelled tissue data may not accurately reflect physiological cfDNA. However, the strong correlations that we found between measured and modelled methylation percentages for individual markers suggests that our data will be robust to such limitations. In addition, we found no differences in the correlation between measured methylation percentage and cell-line content of each sample between the markers that we derived by *in silico* modelling and the markers for breast cancer detection derived from the literature, suggesting again that the markers we developed may have comparable discriminatory utility to the markers developed in other studies.

Although the synthetic cfDNA methylation profiles we created followed the latest knowledge in healthy cfDNA composition (Lam et al. 2017; Moss et al. 2018), the origins of cfDNA in healthy subjects and cancer patients are still not well understood and therefore the composition that we used may not be as accurate as we intend. Further, while the biological noise we introduced here by adding 10% of a mix of six different tissues should make the models somewhat robust to various biological conditions, especially to degenerative or destructive breast lesions, as normal breast is included, only a limited range of tissues was included in our modelling, so even our best CpGs may not be applicable to people with underlying conditions affecting other tissues. Finally, we used regularised linear modelling to derive our *in silico* marker set which led to models with predictive power to detect tumour DNA in cfDNA, with as little as 0.25% of tumour DNA fraction in synthetic cfDNA. As a follow-up to this study, investigating non-linear modelling strategies might lead to additional improvement in model performances, although as the complexity of models increases so does their likelihood of overfitting the data. Opportunities to apply alternative modelling strategies will further expand as the number of publicly available breast tumour samples increases.

The methylation markers that were derived from the literature came from several different sources, each with different approaches to marker development. The study by de Almeida et al. provided 368 CpG markers to distinguish breast tumour from normal breast tissue (de Almeida et al., 2019). They used TCGA methylation data and filtered the CpGs so that only CpGs located within gene bodies and proximal gene regions were included. They ranked their CpGs based on OncoScore to determine cancer-related genes and then used Kaplan-Meier survival curves and log-rank tests to compare the survival curves. They also looked at the correlation between methylation and gene expression from the METABRIC database. This is therefore a well curated gene-centric marker set including CpGs in proximal gene regions. Fackler et al. reported two panels, one of 100 CpGs, and a more selected panel of 30 CpGs in which higher levels of methylation were associated with greater probability of 5-year recurrence following therapy (Fackler et al., 2020). Log-rank and logistic regression models were used to determine the association between DNA methylation and recurrence. These models used a filtered set of beta values (β value ratio ≥1.5) from the 450K data from their samples. It is worth noting that this study was designed to detect recurrence, so it might not have the same power for early detection. Moss et al. focussed on three regions arbitrarily selected from 13 candidate sites identified from TCGA 450K data to show differential methylation between breast cancer samples, other cancer samples and 16 normal tissues (Moss et al. 2020). Only markers with at least 5 CpGs in a 165bp interval were considered as suitable markers. The three markers published in the paper by Moss et al. and the remaining 10 candidate sites are all included in the marker set published here. Uehiro et al. (Uehiro et al., 2016) developed a marker set of 12 CpGs, derived from a screen of breast cancer tissues and cell lines compared to non-cancer tissues and validated in cfDNA from breast cancer patients and healthy volunteers. The four markers showing best detection performance were then validated in an independent set of cfDNA samples.

The 100 markers that we derived by data mining and modelling showed no overlap with the 110 markers that we derived from the literature. However, when the correlation between the *in vitro* methylation data for individual markers that we generated and the 450K data for MCF-7 and leukocytes were compared, we found no difference between the correlations observed for the data mining set and the literature derived set of markers. This suggests that the data mining and literature-derived markers may provide independent yet complementary ability to detect breast cancer DNA in a complex mix such as is encountered in cfDNA.

The experimental method developed here used enzymatic conversion of unmethylated cytosine rather than bisulphite conversion, because enzymatic conversion results in less damage and loss to the input DNA (Vaisvila et al., 2021). Our protocol combines target selection of converted DNA with amplification, deep sequencing and barcoding to achieve more than 500x coverage per sample in target regions, which was sufficient for the spike-in range used here though this could be increased for analysis of cfDNA samples with low or very low ctDNA content. The protocol achieved high conversion rates, with 0.2% methylation called on RepliG-amplified unmethylated control DNA and 96% methylation percentage on HeLa DNA. The high correlations observed between MCF-7 percentage DNA input and measured methylation percentage, and between measured methylation percentage and *in silico* modelled methylation percentage derived from 450K array data indicate the accuracy of the assay and its potential for use in measuring methylation in plasma cfDNA from breast cancer patients.

To date, we have only tested our marker sets for their ability to distinguish leukocyte DNA from MCF-7 DNA and normal breast tissue. While these results show promise, we now aim to test the assay with our set of 210 CpGs on DNA from breast cancer tissue and cfDNA samples from breast cancer patients and healthy subjects to confirm its sensitivity and specificity for detection of breast cancer ctDNA in a direct clinical context.

## Acknowledgements

We thank Nonacus Ltd for a very generous provision of reagents for the experiments as well as many technical discussions in protocol development. We thank Dr Duncan Sproul for advice on the methylation analysis pipeline and for sharing his scripts. We thank Dr Felix Krueger for having extensive data analysis discussions on his tool Bismark. We thank Gil Tomás for his contribution in development of the *in silico* dataset and for valuable discussions. The results described here are in part based upon data generated by the TCGA Research Network: https://www.cancer.gov/tcga.

## Legends for supplementary materials

**Supplementary Table 1**. Genomic coordinates of the 210 CpG markers on the GRCh38 genome reference. Source indicates whether the CpG marker was extracted from the literature (with reference to publications by first author) or from our modelling strategy, as described in the Methods section. The column cg_id refers to the CpG unique identifier associated with each marker.

**Supplementary Table 2**. Linear models selected on synthetic cell-free DNA methylation profiles. We show the area under the ROC curve (AUC) and number of features (non-zero coefficients associated with a single CpG) for each of the 70 models obtained from our *in silico* classification analysis for 7 different percentages or breast tumour DNA and 10 replicates of each.

**Supplementary Table 3**. Total number of fragments sequenced per sample. A fragment (or read pair) is composed of 150 bp paired-end reads. Each fragment is counted only once, with the total number of fragments corresponding to the total number of Read 1 (or Read 2) reads. The composition of each sample is indicated in the ‘Sample content’ column. The ‘Sample names’ column refers to the names associated with each sample.

**Supplementary Table 4**. Mapping statistics per sample. The depth of coverage and total number of mapped reads are indicated for each sample. The composition of each sample is indicated in the ‘Sample content’ column. The ‘Sample names’ column refers to the names associated with each sample.

**Supplementary Figure 1:** Examples of receiver operating characteristic (ROC) curves for models trained for breast cancer detection in synthetic cfDNA under the assumption of 10% random other tissue contribution. For each replicate, a set of 658 synthetic cfDNA samples was assembled, half of which were supplemented with increasing breast cancer profile percentages (0.01%, 0.1%, 0.25%, 0.5%, 0.75%, 1% and 10%). Here we show the ROC curves for replicate 1 only. Also shown are the areas under the ROC curve (AUC) for the predictions of each optimal model, obtained according to the procedure depicted in the Methods section.

**Supplementary Figure 2**. Heatmaps showing Pearson correlation coefficient scores between samples for marker panel CpGs selected to show (A) hypermethylation (Hyper_M) or (B) hypomethylation (Hypo_M) in breast cancer DNA.

