## Supplementary Figure 1 for "Development of methylation-based biomarkers for breast cancer detection by model training and validation in synthetic cell-free DNA"

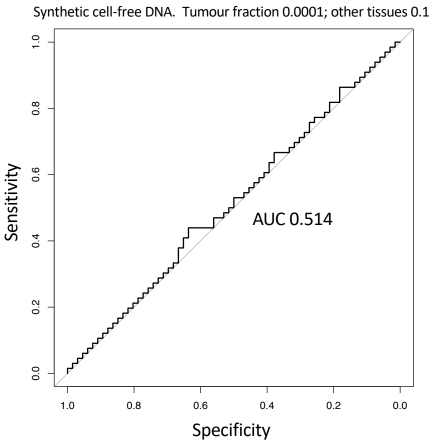


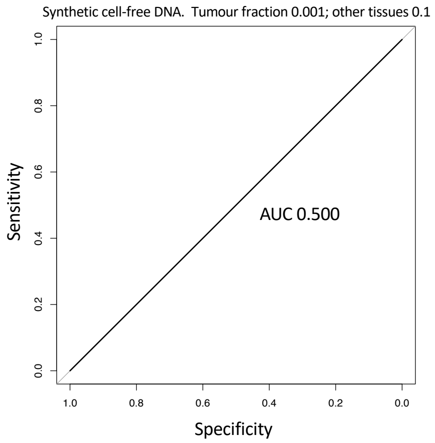


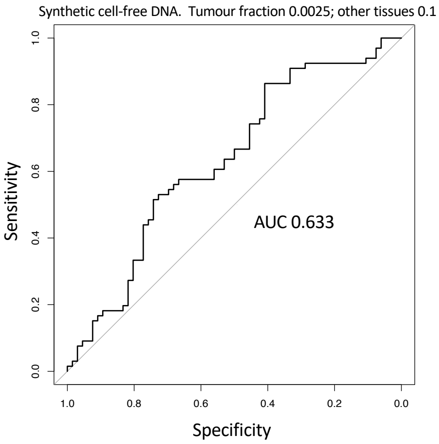


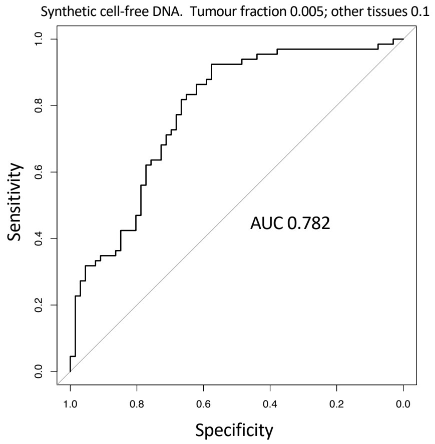


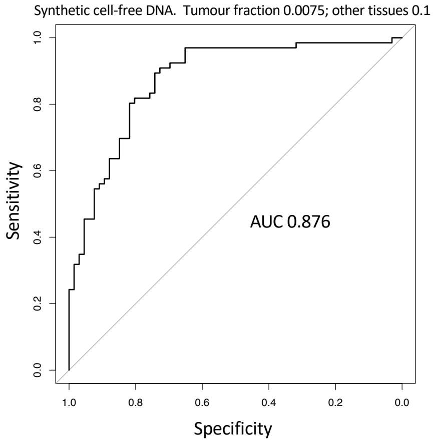


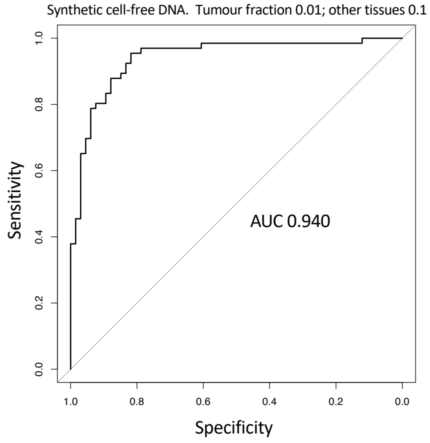


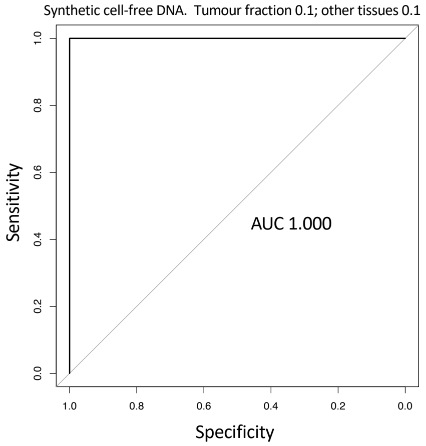


**Supplementary Figure 1:** Examples of receiver operating characteristic (ROC) curves for models trained for breast cancer detection in synthetic cfDNA under the assumption of 10% random other tissue contribution. For each replicate, a set of 658 synthetic cfDNA samples was assembled, half of which were supplemented with increasing breast cancer profile percentages (0.01%, 0.1%, 0.25%, 0.5%, 0.75%, 1% and 10%). Here we show the ROC curves for replicate 1 only. Also shown are the areas under the ROC curve (AUC) for the predictions of each optimal model, obtained according to the procedure depicted in the Methods section.
