## Supplementary Figure 2 for "Development of methylation-based biomarkers for breast cancer detection by model training and validation in synthetic cell-free DNA"

A

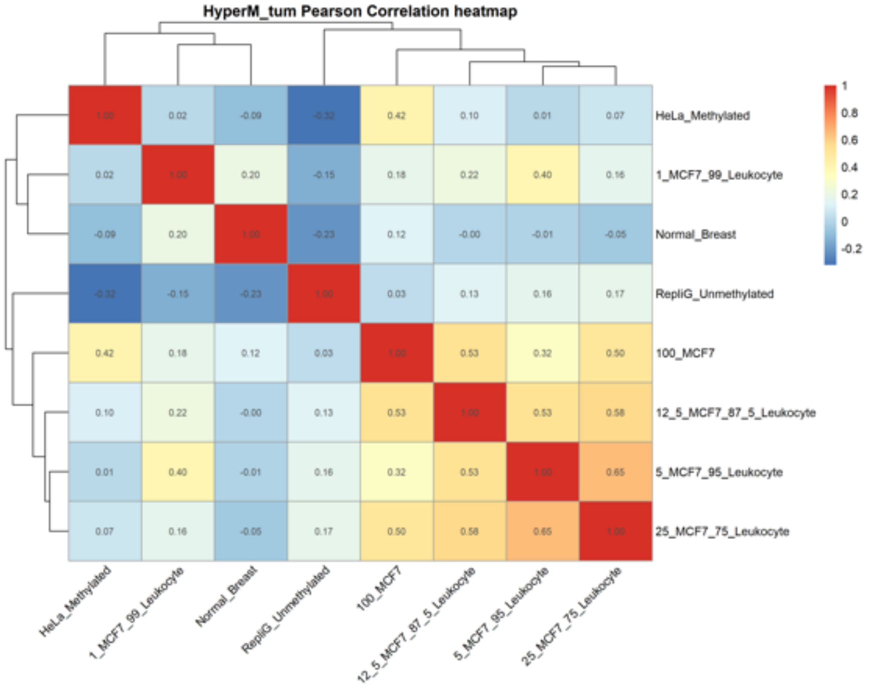


B

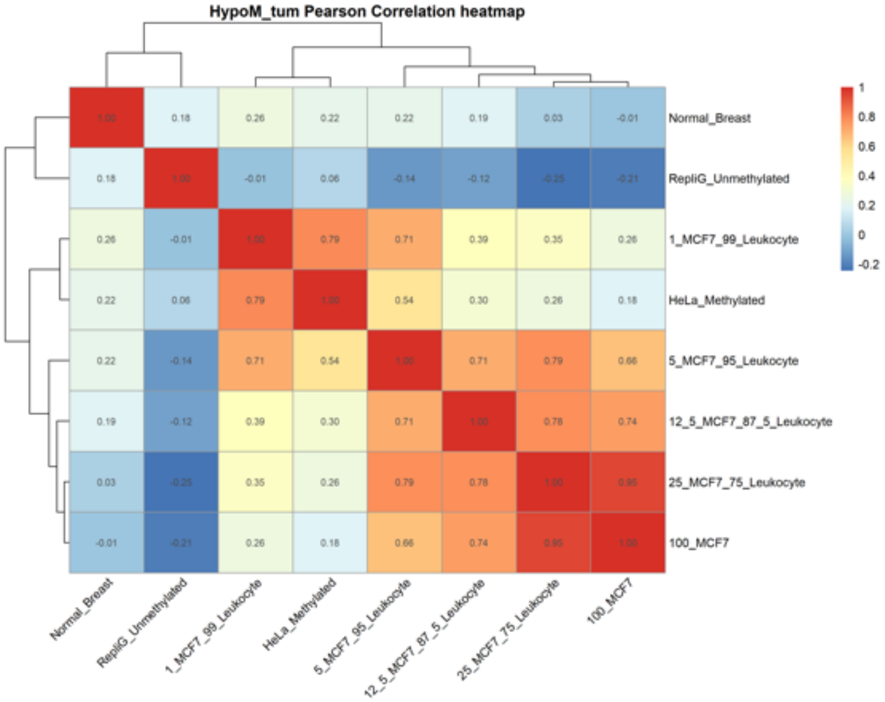


**Supplementary Figure 2.** Heatmaps showing Pearson correlation coefficient scores between samples for marker panel CpGs selected to show (A) hypermethylation (Hyper_M) or (B) hypomethylation (Hypo_M) in breast cancer DNA.
